# Evaluation of the potassium channel tracer [^18^F]3F4AP in rhesus macaques

**DOI:** 10.1101/2020.04.28.065094

**Authors:** Nicolas J. Guehl, Karla M. Ramos-Torres, Clas Linnman, Sung-Hyun Moon, Maeva Dhaynaut, Moses Q. Wilks, Paul K. Han, Chao Ma, Ramesh Neelamegam, Yu-Peng Zhou, Brian Popko, John A. Correia, Daniel S. Reich, Georges El Fakhri, Peter Herscovitch, Marc D. Normandin, Pedro Brugarolas

## Abstract

Demyelination causes slowed or failed neuronal conduction and is a driver of disability in multiple sclerosis and other neurological diseases. Currently, the gold standard for imaging demyelination is MRI, but despite its high spatial resolution and sensitivity to demyelinated lesions, it remains challenging to obtain specific and quantitative measures of demyelination. To understand the contribution of demyelination in different diseases and to assess the efficacy of myelin-repair therapies, it is critical to develop new *in vivo* imaging tools sensitive to changes induced by demyelination. Upon demyelination, axonal K^+^ channels, normally located underneath the myelin sheath, become exposed and increase in expression, causing impaired conduction. Here, we investigate the properties of the K^+^ channel PET tracer [^18^F]3F4AP in primates and its sensitivity to a focal brain injury that occurred three years prior to imaging. [^18^F]3F4AP exhibited favorable properties for brain imaging including high brain penetration, high metabolic stability, high plasma availability, high reproducibility, high specificity, and fast kinetics. [^18^F]3F4AP showed preferential binding in areas of low myelin content as well as in the previously injured area. Sensitivity of [^18^F]3F4AP for the focal brain injury was higher than [^18^F]FDG, [^11^C]PiB and [^11^C]PBR28, and compared favorably to currently used MRI methods.

## INTRODUCTION

Myelin facilitates rapid propagation of action potentials and provides trophic support and protection to axons^1^. As such, the integrity of the myelin sheath is critical for most brain functions including cognition, visual processing and ambulation^2–4^. In addition to multiple sclerosis (MS)^5^, demyelination is emerging as a prominent cause of disability of many brain diseases including spinal cord injury (SCI)^6^, traumatic brain injury (TBI)^7^, stroke^8^ and Alzheimer’s disease (AD)^9^. Currently, demyelination can be detected by histopathology, by PET using tracers that bind to myelin such as the amyloid tracer [^11^C]PiB or [^11^C]MeDAS^10, 11^ or by MRI^12, 13^. Unfortunately, histopathology is not applicable for *in vivo* disease monitoring, myelin tracers are not ideal due to binding to multiple targets and the abundance of myelin, and while structural and diffusion MRI offer some insight into the structural integrity of white matter, MRI measurements are not specific to white matter tissue properties such as myelination or fiber density^14^. Furthermore, MRI offers limited sensitivity to gray matter demyelination^15^. Therefore novel molecular imaging tools for monitoring demyelination *in vivo* are needed.

A well-established biochemical effect of demyelination that is responsible for the poor conduction of action potentials in demyelinated fibers is the dysregulation of axonal voltage-gated K^+^ (K_v_) channels^16^. K_v_ channels are transmembrane proteins that allow the selective passage of K^+^ ions across the cell membrane upon membrane depolarization. In the brain, K_v_ channels are primarily expressed in neurons where they are involved in the propagation of action potentials^17^. In myelinated axons, K_v_1.1 and K_v_1.2 channels are located near the nodes of Ranvier beneath the myelin sheath^18^. Upon demyelination, these protein channels disperse thoughout the demyelinated segment and increase in expression several fold resulting in slowed or failed axonal conduction^16, 19, 20^. 4-aminopyridine (4AP) is a blocker of K_v_1 channels and is used clinically to enhance conduction of demyelinated fibers^21, 22^. This mechanism has been exploited for the symptomatic treatment of MS^23, 24^, SCI^25^ and stroke^26^.

3-fluoro-4-aminopyridine, 3F4AP, is a fluorinated analog of 4AP that binds with similar affinity as 4AP to K_v_1 channels expressed in *Xenopus* oocytes as well as explanted optic nerves from mice^27, 28^. It has been recently shown by autoradiography that [^14^C]4AP and [^14^C]3F4AP accumulate in demyelinated areas in rodent brain after intravenous administration^27^. This is due to the increased level of expression of K_v_1 channels in those areas as well as the increased accessibility of the drug to the channels given the lack of myelin^19, 20, 29, 30^. It has also been reported that 3F4AP can be labeled with fluorine-18^31–33^ and that radiofluorinated [^18^F]3F4AP can be used to detect chemically-induced demyelinated lesions in rat brains using PET^27^. The high sensitivity of this tracer towards demyelination makes it a promising candidate for imaging many neurological diseases where there is demyelination including MS, TBI, SCI, stroke, AD, and others^30^. This tracer also holds promise as a tool for monitoring remyelinating therapies, a major focus in current drug development^30, 34^.

In order for a PET tracer to be of clinical value, it must not only bind to a clinically relevant target, it must also have appropriate pharmacokinetics that allow reliable quantification of the target. Ideally, a brain tracer should have high brain penetration, high metabolic stability, high plasma availability, relatively fast kinetics and high specificity for its target^35^. The goal of this study was to comprehensively evaluate [^18^F]3F4AP in nonhuman primates in preparation for human studies.

## MATERIALS AND METHODS

### Study design

The primary objective was to characterize [^18^F]3F4AP in nonhuman primates. Screening against a panel of 38 other common brain receptors was also performed to rule out off-target binding. [^18^F]3F4AP whole-body scans were performed on two rhesus macaques (Monkey 1 and Monkey 2) to provide a whole-body dosimetry estimation. In two other rhesus macaques (Monkey 3 and Monkey 4), a total of 8 dynamic PET imaging studies with arterial blood sampling were performed to provide a thorough characterization of [^18^F]3F4AP *in vivo* pharmacokinetics in the monkey brain. One of the animals (Monkey 4) showed an incidental finding related to a previously sustained focal brain injury and was also scanned with the well-characterized PET tracers [^11^C]PBR28, [^11^C]PiB and [^18^F]FDG to investigate the nature of the injury. In addition, structural, magnetization transfer and diffusion MRI were performed to assist in brain regions delineation and assess pathological changes.

### Animal studies

All experiments involving animals were performed in accordance with relevant guidelines and regulations and with approval from the institutional animal care and use committees (IACUC) at the Massachusetts General Hospital and NIH. The animals used in this study were adult male monkeys (ages: 9-15 years). Prior to each study, animals were sedated with ketamine/xylazine (10/0.5 mg/kg IM) and were intubated for maintenance anesthesia with isoflurane (1-2% in 100% O_2_). A venous catheter was placed for infusion of the radiotracer and, where applicable, an arterial catheter was placed for sampling of the arterial input function. The animal was then positioned on a heating pad on the bed of the scanner for the duration of the study. Additional procedural details are described in the **Supporting Information (SI)**.

### Receptor panel screen

Samples of non-radiolabeled 4AP and 3F4AP were sent to the psychoactive drug screening program (PDSP) from the NIH/NIMH at the University of North Carolina^36^. Specific details for each assay can be found in the Assay Protocol Book available on the website: https://pdspdb.unc.edu/pdspWeb/

### Radiochemistry

Radiochemical syntheses of [^18^F]3F4AP, [^11^C]PBR28, [^11^C]PiB and [^18^F]FDG were performed as previously described^31, 37–39^.

### Whole-body dynamic PET/CT imaging of rhesus macaques with [^18^F]3F4AP

Two rhesus macaques (Monkey 1 and Monkey 2) were scanned on a Siemens mCT PET/CT scanner for 4h as previously described^27^. Procedural details are provided in the **SI**.

### Human radiation dosimetry estimation

Human organ dosimetry estimates were calculated from whole-body dynamic PET data from Monkey 1 and Monkey 2 using OLINDA/EXM software as described in the **SI**.

### Magnetic Resonance Imaging

Brain MRI was performed on Monkey 3 and Monkey 4 using a 3T Biograph mMR (Siemens Medical Systems). The following sequences were used MEMPRAGE (pre- and post-Gd), T2-FLAIR and DTI. Full details are provided in the **SI**.

### Brain dynamic PET/CT imaging of rhesus macaques with [^18^F]3F4AP

Dynamic PET/CT imaging (2-3h) with arterial blood sampling was performed on Monkey 3 and Monkey 4 using a Discovery MI (GE Healthcare). Each animal had two baseline scans which were separated by one month for Monkey 3 and by one year for Monkey 4. Monkey 4 had four other scans with different doses of unlabeled 3F4AP (0.75, 1.25, 2.5 and 4 mg/kg) co-injected with [^18^F]3F4AP. CT scan was acquired before each PET acquisition for attenuation correction of PET images. See **SI** for additional details.

### Brain PET/CT scans with [^11^C]PiB, [^11^C]PBR28 and [^18^F]FDG

Dynamic PET/CT imaging (90 min) with arterial blood sampling was performed on Monkey 4 after administration of [^11^C]PiB and [^11^C]PBR28. Static PET/CT imaging was performed on Monkey 4 one hour after administration of [^18^F]FDG (6 min scan). Additional details about the procedures and quantification are given in the **SI**.

### Arterial blood sampling

Arterial blood samples of 1-2 mL were drawn every 30 seconds immediately following radiotracer injection and decreased in frequency to every 30 minutes toward the end of the scan. [^18^F]3F4AP metabolism was measured from blood samples acquired at 5, 10, 15, 30, 60, 90, 120 and up to 180 minutes. An additional blood sample of 3 mL was drawn immediately prior to tracer injection in order to measure the plasma free fraction *f*_*p*_ of [^18^F]3F4AP.

### Arterial blood processing and radiometabolite analysis

Radioactivity concentration (in kBq/cc) was measured in whole-blood (WB) and subsequently in plasma (PL) following the centrifugation of WB. Radiometabolite analysis was performed using an automated column switching radioHPLC system^40, 41^. Additional details in the **SI**.

### Plasma free fraction f_p_ determination

Arterial plasma samples of 200µL drawn before radiotracer injection were spiked with 444 kBq of [^18^F]3F4AP. Following a 15 min incubation period, radioactive plasma samples were loaded on ultrafiltration tubes (Millipore Centrifree) and centrifuged at 1500g for 15 min at room temperature. *f*_*p*_ was calculated as the ratio of free ultrafiltrate to plasma concentration and corrected for binding to the ultrafiltration tube membrane.

### Image registration and processing

All PET processing was performed with an in-house developed Matlab software that uses FSL^42^. MR data were processed in native space using FSL. Individual brain MR and PET images were aligned into the MRI NIMH Macaque Template (NMT)^43^ and regional TACs were generated for the occipital cortex, parietal cortex, temporal cortex, frontal cortex, hippocampus, amygdala, striatum, thalamus, white matter and whole cerebellum. Additional details are provided in the **SI**.

### Quantitative analysis of [^18^F]3F4AP brain uptake

Regional TACs were analyzed by compartmental modeling using the metabolite-corrected arterial plasma input function. One-(1T) and two-(2T) tissue compartment model configurations were investigated while fixing the vascular contribution of the WB radioactivity to the PET measurements to 5%. The 2T model was also tested in its reversible and irreversible modes. Estimates of the kinetic parameters were obtained using nonlinear weighted least-squares fitting with the weights defined as the frame durations. The regional total volume of distribution (*V*_*T*_)^44^ was calculated as *K*_*1*_/*k*_*2*_ for a 1T compartment model and as (*K*_1_/*k*_2_)×(1 + *k*_3_/*k*_4_) for a 2T model (see **SI** for details on the kinetic parameters). In addition, Logan and multilinear analysis MA1 graphical methods^45, 46^ for estimation of *V*_*T*_ was preformed. Parametric maps of *V*_*T*_ were generated using the Logan method.

### Blinded analysis

Even though the nature of the study did not allow for fully blinded analysis. The investigators evaluating [^18^F]3F4AP on monkey 4, were unaware of an injury at the time of data acquisition and analysis.

### Statistical analysis

All data are expressed as mean value ± one standard deviation (SD) unless otherwise specified. Agreement between methods was assessed by computing the average measured intraclass correlation coefficient (ICC) among methods or models by use of a two-way mixed-effects model with absolute agreement definition. A *p* value of 0.05 or less was considered statistically significant. All outliers were included in the analysis, and no data were excluded. Additional details are provided in the **SI**.

## RESULTS

### 3F4AP does not bind to other common brain receptors with high affinity

3F4AP has been shown to bind to voltage-gated K^+^ channels (Shaker family) with similar affinity as the MS drug 4AP^27, 28^. Nevertheless, the capacity of this molecule to bind to other brain receptors has not been investigated. For this purpose, we evaluated the binding of 4AP and 3F4AP against 38 common brain targets. The results from this screen are shown in Table 1. These results show no significant binding to 37 of the 38 receptors tested and only low affinity binding for histamine H2 receptors (*K*_i_ ~ 5 µM).

**Table 1.** Percent inhibition at 10 µM concentration (n = 4)

| RECEPTOR | 5-HT<sub>1A</sub> | 5-HT<sub>1B</sub> | 5-HT<sub>1D</sub> | 5-HT<sub>1E</sub> | 5-HT<sub>2A</sub> | 5-HT<sub>2B</sub> | 5-HT<sub>2C</sub> | 5-HT<sub>3</sub> | Alpha<sub>1A</sub> | Alpha<sub>1B</sub> | Alpha<sub>1D</sub> | Alpha<sub>2A</sub> | Alpha<sub>2C</sub> |
| --- | --- | --- | --- | --- | --- | --- | --- | --- | --- | --- | --- | --- | --- |
| 4AP | -1.3 | 10.8 | 1.8 | 6.5 | 7.0 | 1.1 | -0.3 | 5.8 | -13.2 | -9.3 | -0.8 | 60.3 | 16.0 |
| 3F4AP | -12.2 | 17.8 | 2.9 | 7.0 | -3.9 | -6.7 | 5.2 | 5.5 | 12.2 | 1.1 | 7.1 | 26.5 | 9.4 |
| RECEPTOR | Beta<sub>1</sub> | Beta<sub>2</sub> | Beta<sub>3</sub> | BZP rat | D2 | D3 | D4 | DAT | DOR | GABAA | H2 | H3 | H4 |
| 4AP | 15.2 | -5.4 | 37.0 | 28.6 | -14.7 | 7.2 | 7.2 | 13.6 | -5.0 | 8.6 | 53.8 | 0.7 | 1.5 |
| 3F4AP | 17.9 | -4.2 | 10.3 | 29.2 | -9.7 | -1.8 | 16.5 | 42.8 | -5.3 | 8.6 | 55.9 | 5.6 | 32.7 |
| RECEPTOR | HERG | KOR | M1 | M2 | M4 | M5 | MOR | NET | PBR | SERT | Sigma 1 | Sigma 2 |  |
| 4AP | -5.0 | 6.6 | -15.1 | 4.8 | 34.0 | 37.7 | -0.3 | 8.5 | 14.5 | 5.8 | 10.5 | -5.8 |  |
| 3F4AP | -5.3 | 12.3 | -16.1 | 0.9 | 15.6 | 19.6 | -5.1 | -26.1 | 16.5 | 10.5 | 1.3 | 13.4 |  |

### Whole body dynamic imaging shows organ doses within typical levels for ^18^F-labeled tracers

In order to estimate the dose of radiation following administration of [^18^F]3F4AP to human subjects, we used data from whole body 4h dynamic scans in two rhesus monkeys (Monkey 1 and Monkey 2) to calculate the human equivalent doses to the major organs (Table 2). This study showed that effective dose is 21.6 ± 0.6 µSv/MBq, which is comparable to other ^18^F-labeled tracers (*e.g.*, whole body [^18^F]FDG effective dose is ~20.0 µSv/MBq ^47^.

**Table 2.** Estimated human organ radiation doses

| Organ | Monkey 1<br>(μGy/MBq) | Monkey 2<br>(μGy/MBq) | Mean<br>(μGy/MBq) | SD<br>(μGy/MBq) |
| --- | --- | --- | --- | --- |
| Adrenals | 20.0 | 20.6 | 20.3 | 0.4 |
| Brain | 16.1 | 19.4 | 17.8 | 2.3 |
| Galbladder Wall | 22.5 | 25.8 | 24.2 | 2.3 |
| *Small Int | 21.1 | 22.2 | 21.7 | 0.8 |
| Stomach Wall | 19.3 | 29.9 | 24.6 | 7.5 |
| *Upper LI Wall | 21.1 | 20.6 | 20.9 | 0.4 |
| Heart Wall | 17.7 | 19.9 | 18.8 | 1.6 |
| Kidneys | 60.4 | 77.0 | 68.7 | 11.7 |
| Liver | 27.0 | 35.1 | 31.1 | 5.7 |
| Lungs | 12.3 | 13.5 | 12.9 | 0.8 |
| Muscle | 15.2 | 15.2 | 15.2 | 0.0 |
| Pancreas | 20.4 | 21.8 | 21.1 | 1.0 |
| Red Marrow | 32.7 | 21.8 | 27.3 | 7.7 |
| Osteogenic Cells | 46.8 | 41.7 | 44.3 | 3.6 |
| Skin | 11.6 | 11.6 | 11.6 | 0.0 |
| Spleen | 18.4 | 27.0 | 22.7 | 6.1 |
| Testes | 19.9 | 18.2 | 19.1 | 1.2 |
| Thymus | 15.8 | 15.8 | 15.8 | 0.0 |
| Thyroid | 18.7 | 18.0 | 18.4 | 0.5 |
| Bladder Wall | 30.6 | 33.1 | 31.9 | 1.8 |
| Whole body | 18.6 | 18.7 | 18.7 | 0.1 |
| **Effective Dose (μGy/MBq)** | **21.2** | **22** | **21.6** | **0.6** |

\*Estimated from whole-body dose.

### [^18^F]3F4AP displays suitable pharmacokinetic properties for PET imaging

Encouraged by prior findings in rodents and primates^27^, the positive dosimetry results and the apparent low off-target binding, we set out to evaluate the pharmacokinetic properties of [^18^F]3F4AP in rhesus macaques. [^18^F]3F4AP was synthesized with high molar activity (> 37.0 GBq/µmol) and high radiochemical purity (> 98% RCC). The tracer was injected intravenously into two rhesus macaques (Monkey 3 and Monkey 4) and their brains were imaged dynamically for 2-3h while arterial blood was sampled for measuring radioactivity concentration time courses in whole blood (WB) and plasma (PL), tracer metabolism as well as plasma-free fraction (*f*_*p*_).

WB radioactivity time course was consistent across scans and animals (Fig. 1A). WB to PL radioactivity concentration ratio quickly reached a plateau and was close to unity (WB/PL = 1.05 ± 0.01, Fig. 1B). Further analysis of the radioactivity in plasma showed a very high *f*_*p*_ (*f*_*p*_ = 0.92 ± 0.03, range: 0.89-0.99) indicating that [^18^F]3F4AP does not bind to plasma proteins. Radio-HPLC analysis of selected plasma samples revealed very slow metabolic degradation with only a very small fraction of metabolites (Fig. 1C) and a very high proportion of parent compound in plasma (> 90%) even after 2-3 hours post injection (Fig. 1D). Figure 1E shows the average metabolite-corrected arterial plasma curve obtained across scans and animals with the corresponding standard deviation. Taken together, the data in blood indicates high plasma availability, fast plasma clearance and unusually high metabolic stability, which are ideal properties for PET tracers.

**Figure 1:**
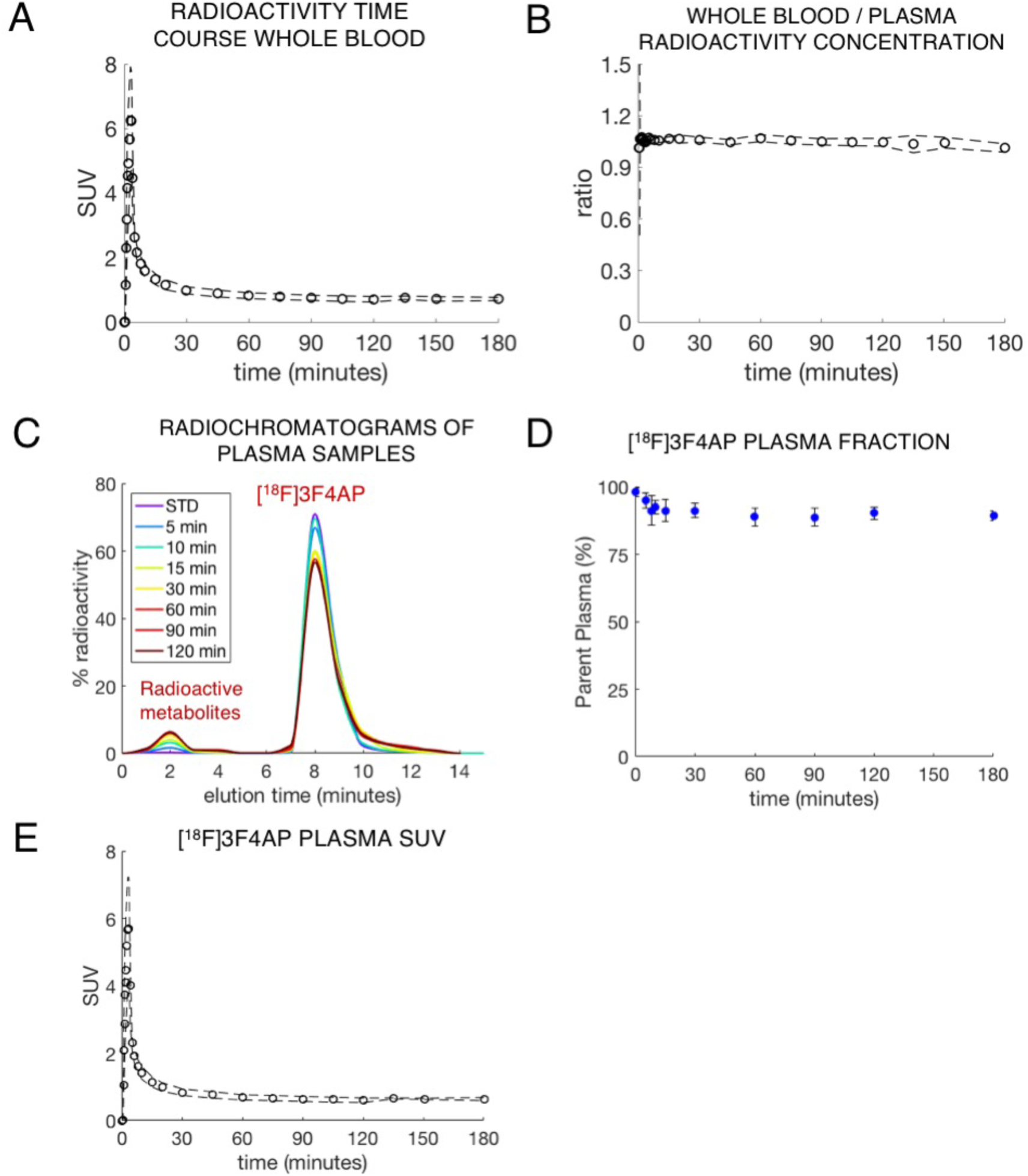
Characterization of [^18^F]3F4AP in blood. (A) Whole-blood SUV time course. (B) Whole-blood to plasma radioactivity concentration ratio. (C) RadioHPLC chromatogram of plasma samples from a representative study. (D) Time course of remaining parent compound in plasma. (E) Metabolite-corrected [^18^F]3F4AP SUV time course in plasma. (Plots A, B, D and E show mean ± s.d. across animals (N=2) and scans (2 in monkey #3 and 6 in monkey#4. Plot C shows data for a representative study).

In the brain, [^18^F]3F4AP peaked quickly (SUV > 3 at ~4 min) and was followed by fast washout. Brain kinetics were fairly homogeneous across brain regions and animals. According to visual inspection of model fits and to the Aikake information criterion (AIC) ^48^, the preferred model (AIC_weight,median_ = 0.999) was a reversible two-tissue compartment model (2T4k) (Fig. 2A and B). *K*_*1*_ values, reflecting tracer delivery, ranged from ~0.34 mL/min/cc in the white matter to ~0.78 mL/min/cc in the striatum, indicating high brain penetration. The *V*_*T*_ estimated from the 2T4k micro-parameters was robust and the time stability of 2T4k *V*_*T*_ estimates was very good as *V*_*T*_ values estimated using only 60 min of data were in excellent agreement with those obtained using 120 min of PET measurements (mean difference = −0.05 ± 0.10 mL/cc, average intraclass correlation coefficient (ICC) = 0.965 with a 95% confidence interval (CI95%) of [0.916, 0.982]). *V*_*T*_ was higher in cortical regions than in the white matter which is consistent with higher expression and accessibility of K^+^ channels in those regions (Fig. 3). *V*_*T*_ values were consistent in both animals and showed low variability across all baseline scans with a coefficient of variation (COV) within 10% for all brain regions surveyed in this work (mean COV = 5.96 ± 2.07%, **Sup. table 1**). Finally, blood-based Logan plots (Fig. 2C and **D**) linearized very well for all datasets by a 30-min *t\** and estimated *V*_*T*_ values were in good agreement with those obtained from the full compartment analysis (2T4k) (mean difference = −0.05 ± 0.07 mL/cc, average measure ICC = 0.976 with CI95% of [0.920, 0.989]). Similar agreement was observed between *V*_*T*_ estimates obtained from MA1 and those obtained from the 2T4k model (mean difference = −0.05 ± 0.07 mL/cc, average measure ICC = 0.973 with CI95% of [0.919,0.987]).

**Figure 2:**
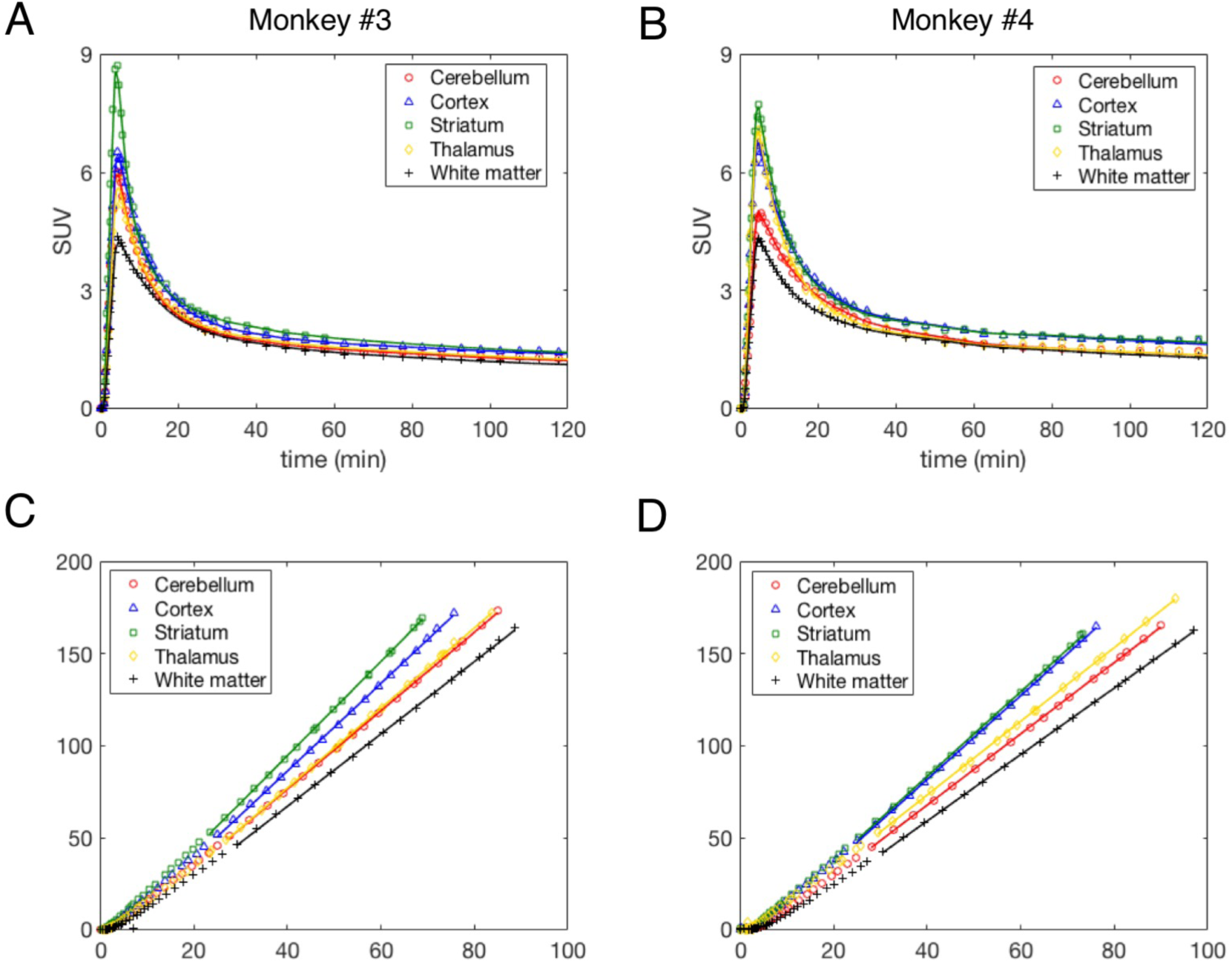
[^18^F]3F4AP kinetics in different brain regions of Monkeys 3 and 4. (A and B) Time-activity curves and 2T4k model fits. (C and D) Logan plots using a 30-min *t\** and corresponding to brain time-activity curves shown in A and B. A and C show data acquired from Monkey 3, while B and D are from Monkey 4. For each monkey, data shown correspond to a single baseline study.

**Figure 3:**
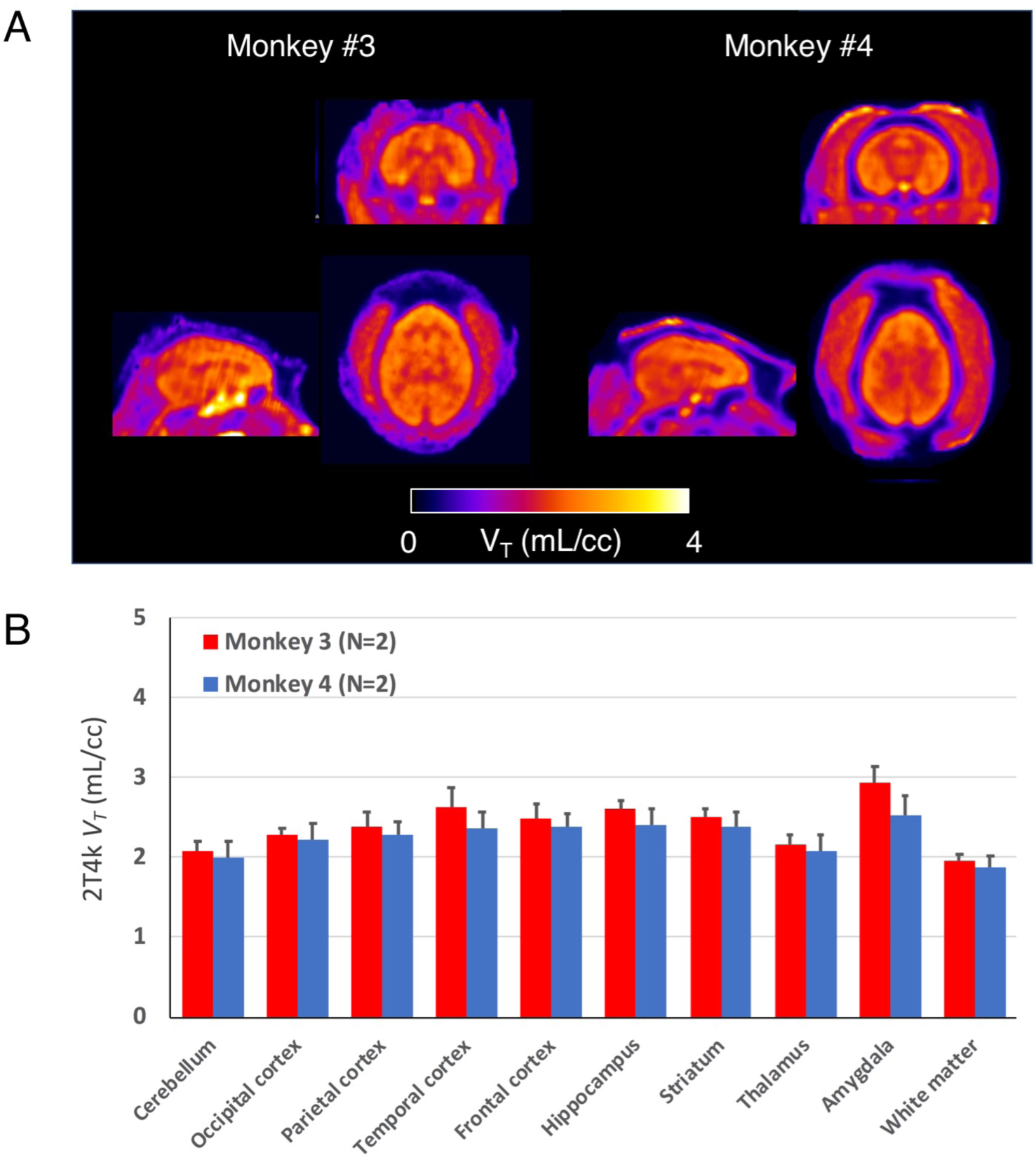
[^18^F]3F4AP in the monkey brain. (A) Representative parametric images of monkey 3 and monkey 4. (B) Mean regional 2T4k *V*_*T*_ and corresponding one SD calculated across studies for each animal.

Finally, motivated by the high metabolic stability of 3F4AP and the stable WB to PL radioactivity concentration ratio, we investigated the use of an image derived input function (IDIF) in lieu of using arterial blood sampling to derive the model input function (IF) for quantification of *V*_*T*_. The IDIF was extracted from a region-of-interest (ROI) positioned in the left ventricular chamber of the heart (**Sup. Fig. 1**). Visually, the obtained image-derived PL curves were in very good agreement with those obtained from arterial blood sampling (**Sup. Fig. 2**) and the area under the curve (AUC) was similar (mean difference = 4.9 ± 0.1%). Consequently, the quantification demonstrated a good agreement in *V*_*T*_ values (mean difference = −0.06 ± 0.07 mL/cc, average measure ICC = 0.963 with CI95% of [0.797, 0.988]).

### It is impractical to block [^18^F]3F4AP signal with unlabeled 3F4AP

Given the low lipophilicity of 3F4AP (logD at pH 7.4 = 0.41) and the negligible off-target binding in brain tissue slices previously reported ^27^, we hypothesized that the [^18^F]3F4AP signal measured in the brain reflects tracer binding to K^+^ channels as well as a nondisplaceable component that mostly reflects free tracer in the intra and extracellular spaces. In order to confirm this hypothesis, we attempted to block the specific signal using unlabeled 3F4AP. For this purpose, we performed four scans in Monkey 4 while coinjecting the tracer with nonradiolabeled 3F4AP at doses ranging from 0.75 to 4 mg/kg. The highest dose tested (4 mg/kg) caused moderate physiological changes including increased heart rate (from 80 to 93 bpm) and increased mean blood pressure (from 55 to 69 mmHg). Furthermore, when the animal arose from anesthesia, it was observed to be shivering, which occurs with K^+^ channel blockers prior to seizures. In mice, 3F4AP as well 4AP caused tremors at ~6 mg/kg (i.p.) and seizures at ~10 mg/kg ^27^. At the doses tested no obvious blocking (assessed by reduction in *V*_*T*_) was observed (**Sup. Table 3**), likely because the doses were below the dose required to saturate the receptors. In fact, an increase in *V*_*T*_ greater than the test/retest variability was observed with increasing doses of 3F4AP which could be due to more channels transitioning to the open state (bindable conformation) upon co-injection of unlabeled 3F4AP (see Discussion).

### [^18^F]3F4AP shows high sensitivity to a focal brain injury

Monkey 4 showed a focal hotspot in a small focal area of the right frontal cortex. The locus of enhanced uptake corresponded to the site of a minor intracranial injury sustained during a craniotomy procedure three years prior to imaging, and was consistent with the burr hole seen on the CT (Fig. 4A). Moreover, the TAC demonstrated very distinct pharmacokinetics with characteristics consistent with increased binding (Fig. 4B). Quantitative analysis showed a 40.3 ± 14.4% (N = 6 scans) higher *V*_*T*_ in the injury site compared to the contralateral site which could not be attributed to changes in perfusion since the quantitative analysis actually demonstrated a lower *K*_*1*_ value (−59.4 ± 10.8%) in this focal spot compared to contralateral site. For comparison, the same analysis was also performed on Monkey 3, which did not undergo a craniotomy and revealed no differences in *V*_*T*_ or *K*_*1*_.

**Figure 4:**
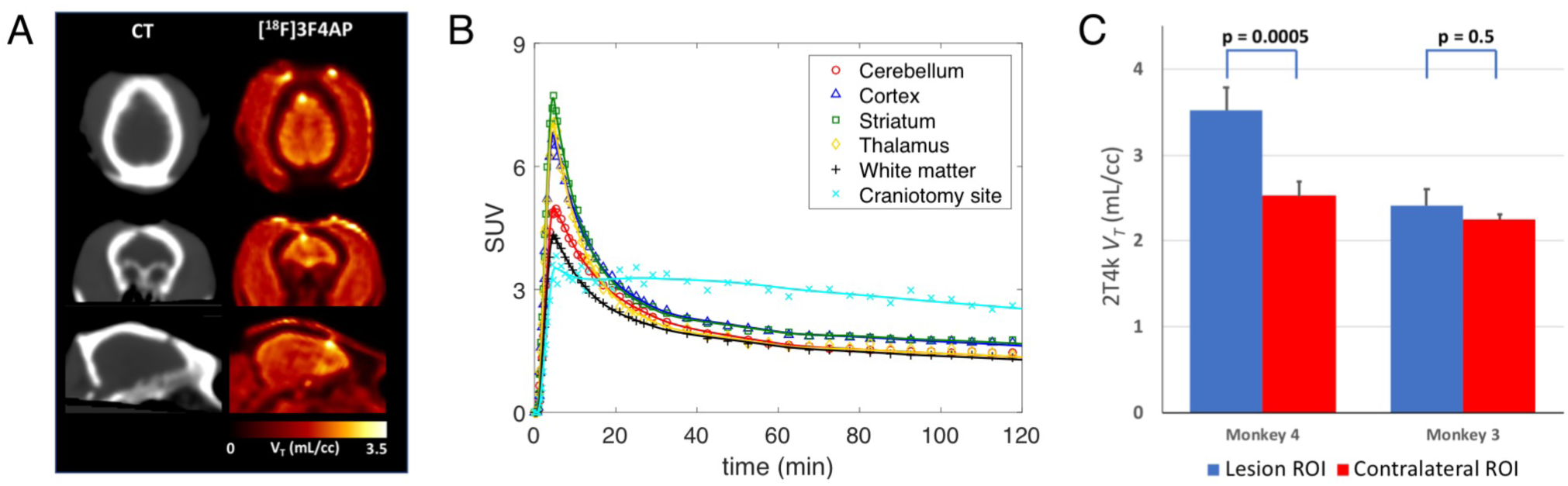
Evaluation of [^18^F]3F4AP in traumatic brain injury. (A) CT and [^18^F]3F4AP images of Monkey 4. (B) Time activity-curve from the lesion (light blue) of Monkey 4 showing very distinct pharmacokinetic as compared to other brain regions. (C) 2T4k *V*_*T*_ in lesion was significantly higher than *V*_*T*_ in contralateral site. For comparison, the other animal showed no differences in corresponding brain regions.

To further investigate the nature of the injury and the potential underlying mechanisms driving [^18^F]3F4AP signal, we performed additional PET scans with [^18^F]FDG, [^11^C]PBR28 and [^11^C]PiB as well as MRI with myelin-specific sequences. [^18^F]FDG was selected to assess tissue metabolism, confirm the presence of living tissue in the affected area and rule out potential processes that could lead to general increase in tracer uptake. [^11^C]PBR28 (TSPO tracer) was selected to assess inflammation^49, 50^, and [^11^C]PIB was selected to assess the presence of amyloid ^51, 52^ as well as demyelination^10, 53^. MRI sequences included T1-weighted multi-echo magnetization-prepared rapid gradient-echo (MEMPRAGE), T2-weighted fluid-attenuated inversion recovery (FLAIR), magnetization transfer ratio (MTR) and diffusion tensor imaging (DTI) as they can inform of demyelination and other pathologies. Images obtained from the different PET tracers and MR sequences are shown in Fig. 5. Qualitatively, [^18^F]3F4AP appeared more sensitive for detecting the lesion than any other imaging methods tested. Quantitatively, ROI-based analysis of the [^18^F]3F4AP images showed the *V*_*T*_ value for lesion to be 5.7 standard deviations higher than the mean *V*_*T*_ of all normal brain regions (Fig. 5B). In comparison, [^18^F]FDG showed a −18% reduction in SUV_60-66min_ in the lesion compared to the contralateral ROI which corresponds to 3.0 standard deviations of the SUV_60-66min_ values of all other brain regions (Fig. 5B). The changes in [^18^F]FDG uptake were suggestive of hypoperfusion and/or hypometabolism ruling out processes that could result in a general increase in tracer binding. In the case of [^11^C]PBR28, the *V*_*T*_ in the lesion was −29.9% lower than the contralateral side, which corresponds to 2.85 standard deviations of the *V*_*T*_ of all other brain regions (Fig. 5B). Furthermore, kinetic analysis of [^11^C]PBR28 showed a −55.1% reduction in *K*_*1*_ in the lesion compared to contralateral ROI. The changes observed with [^11^C]PBR28 indicated reduced perfusion and no discernible inflammation. In the case of [^11^C]PIB, there was a −11.9% decrease in DVR in the lesion compared to the contralateral ROI which corresponds to 1.24 standard deviations of the DVR values of all other brain regions (Fig. 5B). The changes in [^11^C]PIB indicated that there was no amyloid accumulation in the lesion area and possible demyelination.

**Figure 5:**
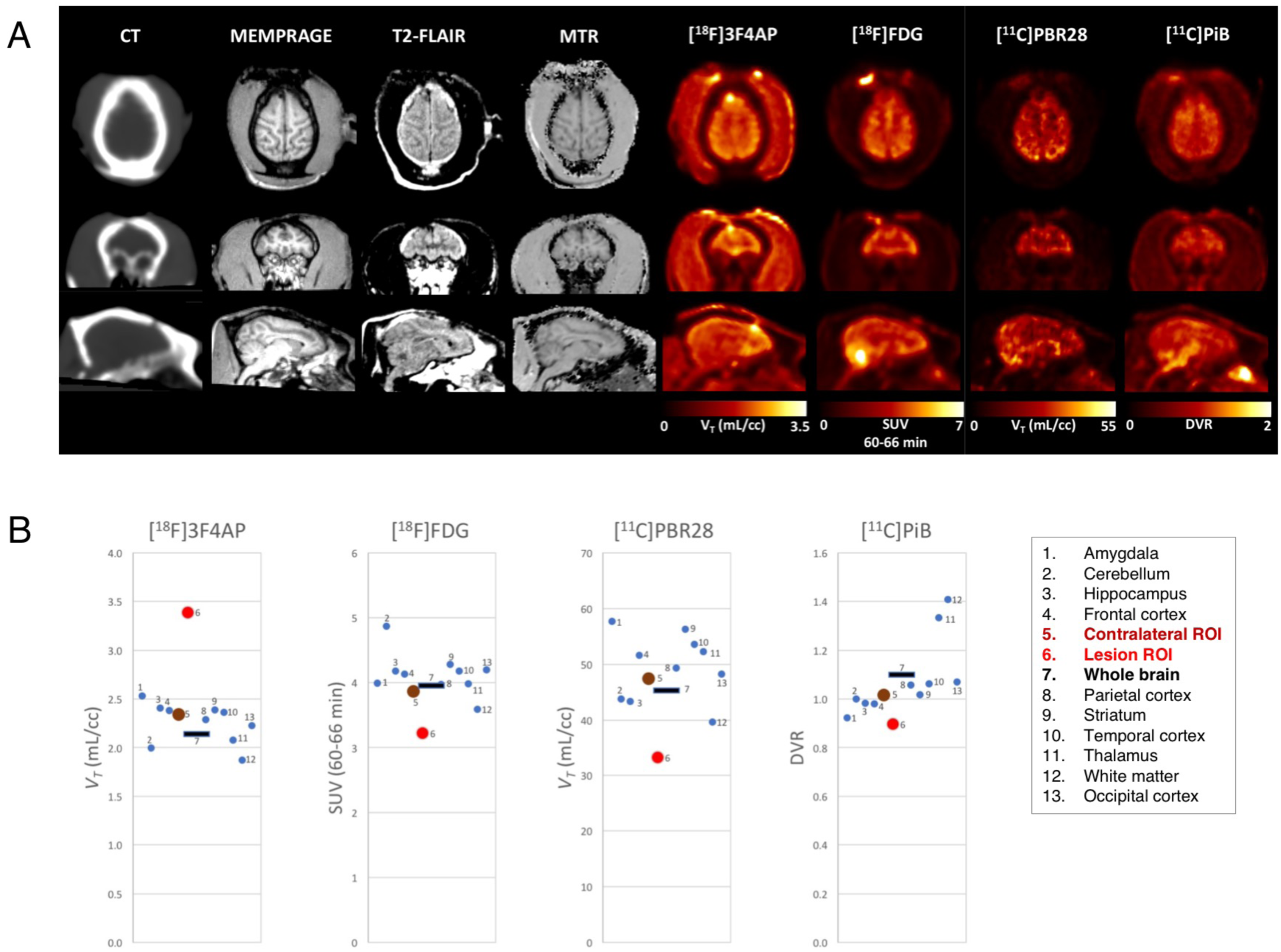
Comparison of [^18^F]3F4AP with other PET radiotracers and imaging modalities. (A) CT, MRI (MEMPRAGE, T2-FLAIR and MTR) and parametric PET images ([^18^F]3F4AP, [^18^F]FDG [^11^C]PBR28 and [^11^C]PiB) of Monkey 4. (B) Brain regional values of the different brain PET tracers.

Results from the quantitative analysis of the MRI data comparing the mean voxel intensity in the lesion to the contralateral mirror ROI are provided in **Supplemental Table 3**. This analysis showed a modest decrease in MEMPRAGE signal (−9% with Cohens *d* = 0.58) as well as T2-FLAIR (−14% with Cohens *d* = 0.58) and MTR (−15% with Cohens *d* = 1.28). There were also small changes on DTI measures that displayed a trend in altered diffusion properties (mean diffusivity and axial diffusivity, *p*=0.10 and *p*=0.05). In addition, no gadolinium contrast enhancement was seen in the lesion. Changes on the MRI were indicative of an intact blood-brain barrier, gliosis and encephalomalacia with possible demyelination. For completeness, the same voxel-based analyses was performed on the PET parametric images ([^18^F]3F4AP, [^11^C]PBR28, [^11^C]PiB and [^18^F]FDG) and the results are also included in **Supplementary Table 3**.

## DISCUSSION

When an area of the brain is injured by an autoimmune attack, a physical blow, a transient lack of oxygen or other insult, one consequence is damage to oligodendrocytes and myelin. Such damage to the myelin sheath hampers the ability of neurons to propagate action potentials, which can cause a myriad of symptoms ranging from cognitive deficits to physical disability. Given that demyelination is potentially reversible and that it is one of the drivers of disability in MS and likely in many other diseases, it is of utmost importance to develop methods to detect demyelination and monitor remyelination.

Currently, demyelination is primarily imaged using MRI. Even though MRI is highly sensitive to demyelinated lesions in the white matter, it lacks specificity as many potentially coexisting pathological processes such as inflammation or axonal loss may result in similar findings. Furthermore, the sensitivity of MRI to gray matter demyelination is low^15^. PET, on the other hand, provides quantitative and biochemically specific information that can complement MRI findings. Previous approaches to image demyelination with PET involve the use of ligands that bind to myelin^10, 11^, which present several limitations including a narrow dynamic range of the measured signal and spill-in signal from adjacent areas due to the limited resolution of PET. Recently, we developed [^18^F]3F4AP, a PET radioligand that binds to voltage-gated K^+^ channels, and demonstrated that it preferentially localizes to demyelinated lesions in several rodent models of demyelination and readily penetrates into the brain of primates ^27^. In view of these promising findings, the primary purpose of the present study was to conduct a thorough evaluation of [^18^F]3F4AP in nonhuman primates including whole-body radiation dosimetry, evaluation of binding to other receptors, and pharmacokinetic modeling.

This present work demonstrates that [^18^F]3F4AP possesses very good properties as a PET radiotracer including negligible binding to other common brain receptors (Table 1), low radiation doses (Table 2), high metabolic stability, low plasma protein binding and suitable kinetics in plasma (Fig. 1) and brain (Fig. 2). [^18^F]3F4AP kinetics in the brain were best described using a reversible two-tissue compartment model with a fixed vascular contribution, which provided a robust quantification of [^18^F]3F4AP signal using the total volume of distribution *V*_*T*_ as the outcome measure. These favorable properties resulted in high repeatability and time stability of *V*_*T*_ estimates as well as low intersubject variability (Fig. 3). In two studies for which we had both the monkey’s heart and brain in the field of view (FOV), we were able to obtain an accurate IDIF from a ROI positioned in the left ventricular chamber (**Sup. Fig. 1** and **2**), which suggests that [^18^F]3F4AP signal may be accurately quantified using standard kinetic modeling methods without the invasive procedure of arterial cannulation.

Our findings also show that it is impractical to block [^18^F]3F4AP signal using unlabeled 3F4AP. Since excessive blockade of K^+^ channels causes seizures^54^, we gradually increased the dose to the highest dose that we estimated would not cause seizures. We had hypothesized that as K^+^ channels become occupied by the unlabeled 3F4AP, there would be a lower number of available channels for the radiotracer to bind and consequently we would observe a reduction in *V*_*T*_. Nevertheless, higher *V*_*T*_ values were observed upon coinjection of unlabeled 3F4AP (**Sup. Table 2**). Upon further examination of this phenomenon, we found a previous report showing that application of high dose of 4AP onto the pial surface of rats caused a local increase in [^3^H]4AP binding due to seizure spreading^55^. This finding leads us to propose a mechanism by which as the concentration of 4AP (or 3F4AP) increases, neuronal firing also increases, which results in more channels in the open, bindable, conformation^56^ leading to an overal increase in binding.

Interestingly, we incidentally observed enhanced [^18^F]3F4AP uptake at the site of a focal intracranial injury sustained during a surgical procedure three years prior to imaging (Fig. 4). We subsequently performed advanced MRI to assess myelin in the region and scanned the animal with tracers for glucose metabolism, [^18^F]FDG, microglial activation, [^11^C]PBR28, and amyloid/myelin, [^11^C]PiB. This comparative study revealed hypoperfusion in the lesion area with no inflammation or amyloid accumulation. Furthermore, the PiB scan was suggestive of demyelination (Fig. 5). From the tracers studied, 3F4AP was the only tracer that showed an increased binding in the injury, and it was also the tracer with the highest apparent sensitivity. On the MRI some of the measures showed clear differences between the lesion and contralateral side (**Sup. Table 3**). For example, MTR – which is commonly interpreted as a marker of myelination showed significant decrease in the lesion. However, other MRI measures such as T2-FLAIR and MEMPRAGE showed changes that are not typically seen in demyelinated lesions. Taken together, MRI findings showed an intact blood-brain barrier, gliosis, encephalomalacia, and possible demyelination. The apparent inconsistency in measures of demyelination may arise from the difficulty of detecting cortical demyelination using conventional MRI^15^ or from upregulation of K^+^ channels through a mechanism independent of demyelination. This is an interesting research question that will be addressed via histopathology when the animal reaches the end of its life.

In summary, this study supports the conclusion that [^18^F]3F4AP is a promising PET radiotracer for imaging voltage-gated K^+^ channels in demyelination, showcases the potential of [^18^F]3F4AP for detecting chronic brain injuries and warrants further investigation in humans. We anticipate that this tracer will be of high clinical utility.

## ACKNOWLEDGMENTS

We thank Daniel Yokell at the MGH Gordon PET Radiopharmacy for providing [^18^F]FDG and [^11^C]PiB, David Lee and Timothy Beaudoin at the MGH Gordon PET Cyclotron for producing ^18^F and ^11^C for the synthesis of [^18^F]3F4AP and [^11^C]PBR28. We thank the veterinary staff (Helen Deng and Eric McDonald) for assistance with animal handling.

## AUTHOR CONTRIBUTIONS

NJG: processed and analyzed the brain PET imaging data and blood data, KMRT: synthesized [^18^F]3F4AP for most of the studies, CL: analyzed the MRI data, MD: processed the blood samples, SHM: performed plasma radioHPLC analysis, MQW: assisted with FDG and MRI scans, PKH and CM: performed the MRI scans, RN: implemented [^18^F]3F4AP synthesis method and synthesized [^11^C]PBR28, YPZ: assisted with the synthesis of [^11^C]PBR28, JAC: performed dosimetry calculations, BP: contributed to the dosimetry study and interpretation of the findings, DSR: contributed to MR interpretation, GEF: contributed to the brain imaging study and data interpretation, PH: performed whole body PET scans, MDN: contributed to the study design, performed monkey brain PET scans and supervised the processing and analysis of PET data, PB: conceived the project, contributed to the study design, the synthesis of [^18^F]3F4AP and supervised the entire project. NJG and PB wrote the manuscript and all authors reviewed and approved it.

## SUPPLEMENTARY INFORMATION AND DATA AVAILABILITY

Supplementary information includes supplemental methods, tables S1-S3 and figures S1, S2 and it is available on the JCBFM website. The datasets generated and/or analysed during the current study are available from the corresponding author on reasonable request.

