## Supplementary Information for "Evaluation of the potassium channel tracer [^18^F]3F4AP in rhesus macaques"

### Contents


| **Topic** | **Display item(s)** | **Page** |
|---|---|---|
| Supplemental Methods | | S3 |
| Supplemental Tables | | S11 |
| **Table S1.** 2T4k regional $V_T$ values measured in Monkey 3 and Monkey 4 for baseline studies. | S1 | S11 |
| **Table S2.** 2T4k regional $V_T$ values measured in Monkey 4 for baseline studies and for studies with co-injection of unlabeled 3F4AP at doses of 0.75, 1.25, 2.5 and 4 mg/kg | S2 | S11 |
| **Table S3.** Comprehensive imaging metrics for the observed lesion and corresponding contralateral ROI. SD is the standard deviation calculated among voxels within the ROI. | S3 | S12 |
| Supplemental Figures | | S13 |
| **Figure S1.** Short axis, horizontal long axis and vertical long axis views of the heart of Monkey 3 | S1 | S13 |
| **Figure S2.** Image derived plasma curves (ID PL) obtained from the left ventricular chamber of Monkey 3 and arterial plasma curves (ART PL) obtained from blood sampling measurements plotted on the same graphs for comparison for baseline scan 1 (A) and baseline scan 2 (B). | S2 | S14 |
| Supplemental References | | S15 |

#### SUPPLEMENTARY METHODS

##### Whole-body dynamic PET/CT imaging of rhesus macaques

Each animal underwent a whole-body (3-bed position) dynamic scan for 4h. CT scan was acquired before the PET acquisition for attenuation correction of PET images. Emission PET data were acquired in three-dimensional (3D) list mode for 240 min following injection of [$^{18}$F]3F4AP. [$^{18}$F]3F4AP was administered via the cubital vein over a 1-minute infusion and was followed by a saline flush. Injected PET tracer activities at the time of injection were 181.3 and 174.3 MBq. Dynamic PET data were reconstructed using a 3D Time-of-Flight OSEM algorithm. Data were framed into dynamic series consisting of 30 frames of 2×15, 4×30, 8×60, 8×120, and 8×240 s per bed position. Final reconstructed images had matrix size of 400×400×411 and voxel sizes of 1.45×1.45×1.45 $mm^3$.

##### Human radiation dosimetry estimation

The main organs were delineated based on the CT and PET images using manual and automated methods. Radioactivity concentration was obtained for each organ at every time point and uncorrected for decay. Non-decay corrected time-activity curves for the major organs and the whole body were extrapolated to 10 half-lives and the area under the curve calculated. OLINDA/EXM Radiation Dose Assessment software was used to estimate the doses to each organ and to the whole body.

##### Magnetic Resonance Imaging

Using a 3T Biograph mMR (Siemens Medical Systems), MRI was performed on Monkey 3 and Monkey 4 for anatomical reference and evaluation of lesion properties on Monkey 4. Data was acquired using the following sequences: a 3D structural T1-weighted multi-echo magnetization-prepared rapid gradient-echo (MEMPRAGE) sequence with repetition time (TR) =2,530 ms, echo time (TE) =1.69 ms, inversion time (TI) =1,100 ms; flip angle = 7°, voxel size = 1×1×1 $mm^3$, matrix size = 256×256×176, and number of averages= 4; a T2-weighted fluid-attenuated inversion recovery (FLAIR) sequence with TR/TE = 11,230/95 ms, TI = 2,500 ms, flip angle = 150°, voxel size = 0.86×0.86×1.4 $mm^3$, and matrix size 256×256×80; a 3D fast low-angle shot (FLASH) sequence with TR/TE=34/6.15 ms, flip angle = 10°, voxel size = 0.5×0.5×2 $mm^3$, and matrix size = 384×512×60, with and without magnetisation transfer saturation; and a diffusion-weighted imaging sequence with 64 directions, TR/TE =6,400/110 ms, flip angle = 90°, voxel size = 2×2×2 $mm^3$, and matrix size = 110×110×38. The T1-weighted imaging was performed using the MEMPRAGE sequence pre- and post-gadolinium injection.

**Positron Emission Tomography imaging of rhesus macaques with [$^{18}$F]3F4AP**

*Imaging procedures:* Monkey 3 and Monkey 4 were scanned on a Discovery MI (GE Healthcare) PET/CT scanner. Each animal had two baseline scans which were separated by one month for Monkey 3 and by one year for Monkey 4. Monkey 4 had four other scans with different doses of unlabeled 3F4AP (0.75, 1.25, 2.5 and 4 mg/kg) co-injected with [$^{18}$F]3F4AP. CT scan was acquired before each PET acquisition for attenuation correction of PET images. Emission PET data were acquired in 3D list mode for at least 120 min and up to 180 minutes following injection of [$^{18}$F]3F4AP. [$^{18}$F]3F4AP was administered via the lateral saphenous vein over a 3-minute infusion and was followed by a 3-minute infusion of saline flush. All injections were performed using syringe pumps (Medfusion 3500). Arterial blood sampling was performed during all dynamic PET acquisition (see next section below). Injected PET tracer activities at the time of injection were 233.5±17.4 MBq (range 211.5 – 258.0). Molar activity ($A_m$) of [$^{18}$F]3F4AP at the time of injection (tracer only) was 72.5 ± 36.7 GBq/µmol (range 30.9 – 120.3, n=4 measurements). Corresponding injected mass was 449.6 ± 236.6 ng (range 206.1 – 773.4, n=4 measurements).

In Monkey 4, the [$^{18}$F]3F4AP scans revealed an incidental finding related to focal brain injury sustained three years prior to imaging (see Results section). In order to further investigate the nature of the injury, this monkey also underwent dynamic scans with [$^{11}$C]PiB and [$^{11}$C]PBR28, as well as a static [$^{18}$F]FDG scan. Details about [$^{11}$C]PBR28, [$^{11}$C]PiB and [$^{18}$F]FDG PET imaging and quantification are given in the next section.

Dynamic PET data were reconstructed using a fully 3D time-of-flight iterative reconstruction algorithm using 3 iterations and 34 subsets while applying corrections for scatter, attenuation, deadtime, random coincident events, detector normalization and incorporating point spread function measurements. For all dynamic scans, list mode data were framed into dynamic series of 6×10 sec, 8×15 sec, 6×30 sec, 8×60 sec, 8×120 sec and remaining were 300 sec frames. Final reconstructed images had voxel dimensions of 256×256×89 and voxel sizes of 1.17×1.17×2.8 $mm^3$.

*Arterial blood sampling*: Arterial blood samples of 1 to 2 mL were drawn every 30 seconds immediately following radiotracer injection and decreased in frequency to every 30 minutes toward the end of the scan. [$^{18}$F]3F4AP metabolism was measured from blood samples acquired at 5, 10, 15, 30, 60, 90, 120 and up to 180 minutes. An additional blood sample of 3 mL was drawn immediately prior to tracer injection in order to measure the plasma free fraction $f_p$ of [$^{18}$F]3F4AP (see below).

*Arterial blood processing:* Radioactivity concentration (in kBq/cc) was measured in whole-blood (WB) and subsequently in plasma (PL) following the centrifugation of WB. Radiometabolite analysis was performed using an automated column switching radio-HPLC (High Performance Liquid Chromatography) system [1, 2]. Briefly, arterial plasma was injected onto the column switching radio-HPLC and initially trapped on a catch column (Waters Oasis HLB 30 µm) using mobile phase consisting of 99:1 10 mM ammonium bicarbonate pH 8 in water:MeCN at 1.8 mL/min (Waters 515 pump). After 4 minutes, the catch column was backflushed with 95:5 10 mM ammonium bicarbonate pH 8 in water:MeCN at 1 mL/min (second Waters 515 pump) and directed onto a Waters XBridge BEH C18 (130 Å, 3.5 µm, 4.6 mm x 100 mm) analytical column. Eluent was collected in 1 minute intervals and subsequently assayed for radioactivity using a Wallac Wizard 2480 gamma counter. After background radioactivity subtraction, radioactivity eluting in the [$^{18}$F]3F4AP peak was divided by the total activity and multiplied by 100% for calculation of percent parent in plasma (%PP). Standardized uptake value (SUV) time courses of radioactivity in WB and in PL were generated by correcting absolute radioactivity concentrations (C [kBq/ml]) for subject body weight (BW [kg]) and injected dose (ID [MBq]): SUV=C/(ID/BW). %PP time course for both monkeys was fit to a single exponential decay plus a constant. Arterial input functions of [$^{18}$F]3F4AP radioactivity concentration in PL $\boldsymbol{C_P(t)}$ were generated by correcting the total PL radioactivity concentration $\boldsymbol{C_P^{total}(t)}$ for radiotracer metabolism using individual %PP(t): $\boldsymbol{C_P(t) = C_P^{total}(t)} \times$

$\%\boldsymbol{PP}(\boldsymbol{t})/100\%$. [$^{18}$F]3F4AP metabolite-corrected arterial SUV time courses were generated. In addition, the plasma free fraction $f_p$ of [$^{18}$F]3F4AP was measured in triplicate by ultrafiltration as follows. Arterial plasma samples of 200 μL drawn before radiotracer injection were spiked with 444 kBq of [$^{18}$F]3F4AP. Following a 15 min incubation period, radioactive plasma samples were loaded on ultrafiltration tubes (Millipore Centrifree) and centrifuged at 1500g for 15 min at room temperature. $f_p$ was calculated as the ratio of free ultrafiltrate to plasma concentration and corrected for binding to the ultrafiltration tube membrane.

*Image registration and processing*: All PET processing was performed with an in-house developed Matlab software that uses FSL [3] for registration purposes. Individual brain MR and PET images were aligned into the MRI National Institute of Mental Health Macaque Template (NMT) [4] according to the following procedure. An early summed PET image (0-10 min post tracer injection) was rigidly co-registered to the animal's individual MEMPRAGE which was subsequently registered into the NMT template space using a 12-parameter affine transformation followed by non-linear warping. The transformation matrices were then combined and applied inversely on the rhesus atlases in order to warp all atlases into the native PET image space for extraction of TACs. Details on the atlases used in this work are provided below. Regional TACs were generated for the occipital cortex, parietal cortex, temporal cortex, frontal cortex, hippocampus, amygdala, striatum, thalamus, white matter and whole cerebellum.

MR data were processed in native space using FSL. After co-registration and alignment, signal from pre- and post-contrast T1-weighted images, T2-weighted image, ratio between T1- and T2-weighted images (T1/T2), and MTR were calculated. Diffusion data was processed using FMRIB´s Diffusion Toolbox [5-7] with eddy current correction, and fractional anisotrophy (FA), mean diffusivity (MD), mode of anisotropy (MO) and radial diffusivity (RD) were calculated in Monkey 4. All calculations were performed for the lesion and the corresponding contralateral region for analysis.

*Anatomical atlases for ROI-based analysis:* Two sets of digital anatomical atlases were aligned to the NMT template space and used for ROI-based analysis and TACs extraction in this work. These were composite ROIs derived from the *Paxinos et al.* rhesus brain regions [8] previously aligned to the *McLaren et al.* rhesus MRI brain template [9] by Moirano et al. as well as composite ROIs derived from the INIA19 NeuroMaps rhesus atlas [10]. Briefly, the T1-weighted average MR images of each template were transformed to the NMT template using a 12-parameter affine registration and non-linear warping both weighted on the brain (i.e. after skull stripping) and the transformation matrices were applied to the composite ROIs.

*Assessment of Image derived input function (IDIF):* In the two studies performed in Monkey 3, we tested the possibility of using an IDIF in lieu of arterial blood measurements. The animal was positioned such as dynamic images were acquired with both the brain and the heart in the field of view (FOV). The dynamic images were reoriented to display the heart in the standardized short axis view [11] and a time activity curve (TAC) was extracted from the left ventricular (LV) chamber using an ellipsoidal ROI positioned in the basal portion (length of each minor axis=8mm, length of major axis=20 mm) (**Fig. S1**). This LV TAC was then corrected for the WB to PL radioactivity concentration ratio using a fix nominal value calculated as the mean WB/PL ratio derived from all studies (WB/PL = 1.05, **Fig. S2)**. The obtained PL curve was then corrected for radiotracer metabolism using individual parent in plasma time course to derive the IDIF. The image-derived PL curves were compared to the arterial plasma curves obtained from blood sampling by visual assessment and by calculating the area under the PL curves. Measurements of $V_T$, the main outcome of interest in this work, obtained while using the IDIF were also compared to those obtained using the IF derived from arterial blood sampling.

**Brain imaging studies with [$^{11}$C]PBR28, [$^{11}$C]PiB and [$^{18}$F]FDG**

Baseline [$^{11}$C]PBR28, [$^{11}$C]PiB, and [$^{18}$F]FDG scans were acquired in Monkey 4 on the same PET/CT scanner as for [$^{18}$F]3F4AP studies and following a similar experimental protocol. Images were reconstructed using the same iterative reconstruction algorithm and parameters. Image registration and processing were performed as described above for [$^{18}$F]3F4AP.

*[[11]C]PBR28 imaging*: Injected PET tracer activity at the time of injection was 179.5 MBq and molar activity ($A_m$) was 92.8 GBq/μmol. Total injected mass was 672.0 ng. Dynamic PET data were acquired for 120 min upon tracer injection. Acquired list mode data were framed into dynamic series of 6x10, 8x15, 6x30, 8x60, 8x120 and 18x300-s frames. Arterial blood sampling was performed during the dynamic PET acquisition because of the absence of a reference region devoid of specific binding that could be used to quantify brain uptake. Arterial blood sampling scheme was similar as for [[18]F]3F4AP studies and metabolism was measured from blood samples acquired at 3, 5, 8, 15, 30, 60, 90, 120 minutes. Radiometabolite analysis was performed using our automated column switching radio-HPLC system. The 99:1 $H_2O$:MeCN mobile phase was used to trap the sample on the catch column during the first four minutes (flow rate = 1.8 mL/min) and a mobile phase of 60:40 0.1 M ammonium formate in water:MeCN was used to backflush the catch column and direct the sample onto the analytical column (flow rate = 1 mL/min). Eluent was collected in 1-minute intervals and subsequently assayed for radioactivity using a Wallac Wizard 1470 gamma counter.

[[11]C]PBR28 signal was quantified using a one-tissue compartment model ($K_1$-$k_2$), as in Imaizumi et al. [12], with the vascular contribution of radioactivity in WB to the PET signal fixed to 5% (1T2kv). The regional total volume of distribution ($V_T$) [13] was calculated as $K_1/k_2$. Parametric map of $V_T$ was calculated using the Logan graphical method with a 65-min $t^*$ [14].

*[[11]C]PiB imaging*: Injected PET tracer activity at the time of injection was 198.1 MBq and molar activity ($A_m$) was 70.0 GBq/μmol. Total injected mass was 725.2 ng. Dynamic PET data were acquired for 120 min upon tracer injection. Acquired list mode data were framed into dynamic series of 6x10, 8x15, 6x30, 8x60, 8x120s and 18x300-s frames. [[11]C]PiB signal was quantified using Logan Distribution Volume Ratio (DVR) [15] and a 30-min $t^*$. The relative delivery $R_1$ was calculated using the simplified reference tissue model (SRTM) [16]. The cerebellar gray matter was used as a reference region (i.e. devoid of specific binding).

*[$^{18}$F]FDG imaging*: Injected PET tracer activity at the time of injection was 438.7 MBq. Static PET images were acquired at one-hour post-tracer injection for 6 min. [$^{18}$F]FDG PET signal was measured using Standard Uptake Value (SUV) and SUV map was generated. SUV is a semiquantitative measure of the tracer uptake that normalizes the measured tracer activity in the image to the injected activity and total body weight and was calculated as SUV= [$C_{PET}$ x $BW$ x 1000]/$ID$, where $C_{PET}$ is the measured tracer concentration in PET image in Bq/cc, $BW$ is the body weight in kg and $ID$ is the injected dose in Bq at the time of imaging.

**Description of kinetic parameters and outcome measure for signal quantification**

For quantification using compartment models describing reversible tracer kinetics, we followed the consensus nomenclature for imaging of reversibly binding radiotracers [13]. Our primary outcome of interest was the macro-parameter $V_T$ which represents the total volume of distribution (the equilibrium ratio of tracer in tissue relative to plasma which is linearly related to tracer binding to the target). In the one-tissue compartment model, the total volume of distribution is calculated as $V_T = K_1/k_2$ where $K_1$ is the rate constant for transfer of tracer from arterial plasma to tissue and $k_2$ represents the efflux rate constant from tissue to plasma. Conversely, in the two-tissue compartment model, the total volume of distribution is calculated as $V_T = K_1/k_2(1 + k_3/k_4)$ where $K_1$ is the rate of transfer from plasma to the non-displaceable compartment in tissue, $k_2$ represents the efflux from the non-displaceable compartment back to plasma, $k_3$ is the rate constant describing tracer binding from the non-displaceable compartment to the specifically bound compartment, and $k_4$ the rate constant representing the unbinding process from the specific compartment back to the non-displaceable compartment. The microparameter $K_1$ is in unit of mL/min/cc while $k_2$, $k_3$ and $k_4$ have unit of min$^{-1}$. Alternatively, the parameter $k_2$ may represent the efflux rate constant from a free tissue compartment to plasma, while $k_3$ may represent binding to a non-specific binding compartment and $k_4$ unbinding from nonspecifically bound to free compartment. It should be noted that such situation is mathematically indistinguishable from the aforementioned case, where $k_3$ and $k_4$ represent specific binding and unbinding processes. For quantification using the Logan graphical analysis method with reference region input function [15], the Distribution Volume Ratio (DVR) is directly estimated and represents the ratio of total volume distribution in the target region and total volume distribution in the reference region [DVR = $V_T$(target)/$V_T$(reference_region)]. The relative delivery $R_1$ represents the ratio of rate constants $K_1$ for transfer of tracer from arterial plasma to tissue in the target region and reference region [$R_1$ = $K_1$(target)/$K_1$(reference_region)].

**Statistical analysis**

All data are expressed as mean value ± one standard deviation (SD) unless otherwise specified. Agreement between methods was assessed by computing the average measured intraclass correlation coefficient (ICC) among methods or models by use of a two-way mixed-effects model with absolute agreement definition. ICC values were presented along with the 95% confidence interval (CI95%). The AIC was used to assess the relative goodness of fit between compartment models [17]. AIC weights [18] were computed to evaluate the probability of one model being preferred over the other candidates. The coefficient of determination $R^2$ and $t$ distribution of the Fisher transformation were used to generate $p$ values for linear regressions and to assess correlation between $V_T$ and $K_1$. A paired t-test was used to compare the mean $V_T$ values across scans between lesion and contralateral side. For analyses related to Table 4 summarizing the MR imaging findings and corresponding metrics, unpaired t-test were used to calculate $t$ and $p$ values and Cohens's $d$ effect size was calculated. A $p$ value of 0.05 or less was considered statistically significant. All outliers were included in the analysis, and no data were excluded.

#### SUPPLEMENTARY TABLES

**Table S1**. 2T4k regional $V_T$ values measured in Monkey 3 and Monkey 4 for baseline studies.

| ROI | Monkey 3 | | Monkey 4 | | mean | SD | COV (%) |
|---|---|---|---|---|---|---|---|
| | Baseline 1 | Baseline 2 | Baseline 1 | Baseline 2 | | | |
| Cerebellum | 2.17 | 2.00 | 1.94 | 2.05 | 2.04 | 0.10 | 4.73 |
| Occipital cortex | 2.34 | 2.23 | 2.21 | 2.24 | 2.26 | 0.06 | 2.52 |
| Parietal cortex | 2.52 | 2.25 | 2.27 | 2.31 | 2.34 | 0.13 | 5.39 |
| Temporal cortex | 2.80 | 2.45 | 2.31 | 2.42 | 2.49 | 0.21 | 8.55 |
| Frontal cortex | 2.62 | 2.35 | 2.36 | 2.40 | 2.43 | 0.13 | 5.18 |
| Hippocampus | 2.68 | 2.56 | 2.23 | 2.59 | 2.51 | 0.20 | 7.83 |
| Striatum | 2.58 | 2.42 | 2.29 | 2.50 | 2.45 | 0.13 | 5.14 |
| Thalamus | 2.24 | 2.07 | 1.95 | 2.21 | 2.12 | 0.14 | 6.45 |
| Amygdala | 3.08 | 2.80 | 2.48 | 2.59 | 2.74 | 0.27 | 9.71 |
| White matter | 2.01 | 1.89 | 1.82 | 1.93 | 1.91 | 0.08 | 4.07 |
| Whole brain | 2.29 | 2.16 | 2.10 | 2.18 | 2.18 | 0.08 | 3.63 |
| | | | | | | Mean | 5.96 |
| | | | | | | St.dev. | 2.07 |

**Table S2**. 2T4k regional $V_T$ values measured in Monkey 4 for baseline studies and for studies with co-injection of unlabeled 3F4AP at doses of 0.75, 1.25, 2.5 and 4 mg/kg.

| ROI | Baseline 1 | Baseline 2 | 0.75 mg/kg | 1.25 mg/kg | 2.5 mg/kg | 4 mg/kg |
|---|---|---|---|---|---|---|
| Cerebellum | 1.94 | 2.05 | 2.04 | 2.44 | 2.14 | 2.37 |
| Occipital cortex | 2.21 | 2.24 | 2.27 | 2.70 | 2.40 | 2.52 |
| Parietal cortex | 2.27 | 2.31 | 2.29 | 2.66 | 2.45 | 2.56 |
| Temporal cortex | 2.31 | 2.42 | 2.45 | 2.90 | 2.62 | 2.66 |
| Frontal cortex | 2.36 | 2.40 | 2.44 | 2.79 | 2.47 | 2.60 |
| Hippocampus | 2.23 | 2.59 | 2.39 | 2.79 | 2.57 | 2.73 |
| Striatum | 2.29 | 2.50 | 2.34 | 2.79 | 2.44 | 2.57 |
| Thalamus | 1.95 | 2.21 | 2.08 | 2.49 | 2.17 | 2.46 |
| Amygdala | 2.48 | 2.59 | 2.67 | 3.10 | 2.77 | 2.95 |
| White matter | 1.82 | 1.93 | 1.87 | 2.19 | 1.97 | 2.15 |
| Whole brain | 2.10 | 2.18 | 2.18 | 2.53 | 2.28 | 2.42 |

**Table S3**. Comprehensive imaging metrics for the observed lesion and corresponding contralateral ROI. SD is the standard deviation calculated among voxels within the ROI.

| Scan | Lesion (SD) | Contralateral (SD) | *t* | *p* | Signal change (%) | Cohens *d* |
|---|---|---|---|---|---|---|
| $T1_{Pre}$ | 286 (62.8) | 315 (33.3) | 4.3 | <0.001 | -9.0 | 0.6 |
| $T1_{Post}$ | 321 (45.7) | 322 (35.7) | 0.2 | n.s | 0.3 | 0.0 |
| $T1_{Post-Pre}$ | 35.1 (38.1) | 7.03 (34.3) | 6.3 | <0.0001 | 400.0 | 0.8 |
| T2 | 339 (47.3) | 395 (29.2) | 11.6 | <0.0001 | -14.0 | 1.4 |
| T1/T2 | 0.854 (0.209) | 0.804 (0.126) | 2.3 | 0.02 | 6.2 | 0.3 |
| MTR | 0.359 (0.0614) | 0.422 (0.0333) | 10.4 | <0.0001 | -15.0 | 1.3 |
| FA | 0.197 (0.0578) | 0.209 (0.0920) | 0.4 | 0.6847 | -5.7 | 0.2 |
| MD | 0.000599 (0.000142) | 0.000668 (0.000062) | 1.7 | 0.1077 | -10.3 | 0.6 |
| MO | 0.244 (0.594) | 0.607(0.442) | 1.8 | 0.0775 | -59.9 | 0.7 |
| L1 | 0.000718(0.000166) | 0.000823 (0.000095) | 2.1 | 0.0501 | -12.8 | 0.8 |
| RD | 0.000539 (0.000135) | 0.00059 (0.000072) | 1.2 | 0.2234 | -8.6 | 0.5 |
| **$[^{18}F]$3F4AP** | **3.13 (0.29)** | **2.29 (0.18)** | 15.6 | **<0.0001** | **37.0** | **3.4** |
| $[^{11}C]$PBR28 | 17.04 (5.21) | 23.60 (4.65) | 10.8 | <0.0001 | -28.0 | 1.3 |
| $[^{11}C]$PiB | 0.87 (0.16) | 0.99 (0.12) | 6.9 | <0.0001 | -12.0 | 0.8 |
| $[^{18}F]$FDG | 3.29 (0.39) | 3.92 (0.40) | 12.9 | <0.0001 | -16.0 | 1.6 |

MTR magnetization transfer ratio, FA fractional anisotropy, MD mean diffusivity, MO mode of anisotropy, L1 axial diffusivity, RD radial diffusivity

**SUPPLEMENTARY FIGURES**

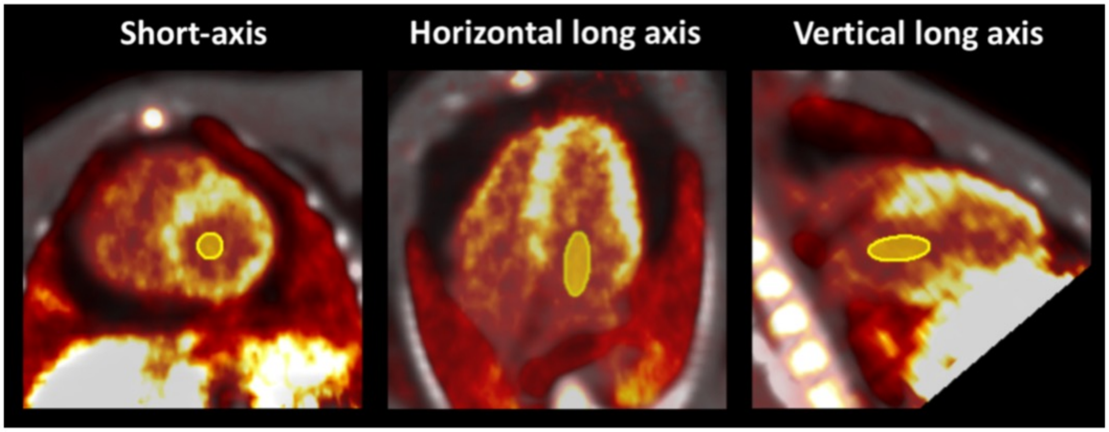


**Figure S1: Short axis, horizontal long axis and vertical long axis views of the heart of Monkey 3.** Summed PET images of early frames (4-10 minutes) are shown with superimposed CT images for anatomical information. ROI for extraction of the TAC in the basal portion of the LV chamber is shown in yellow.

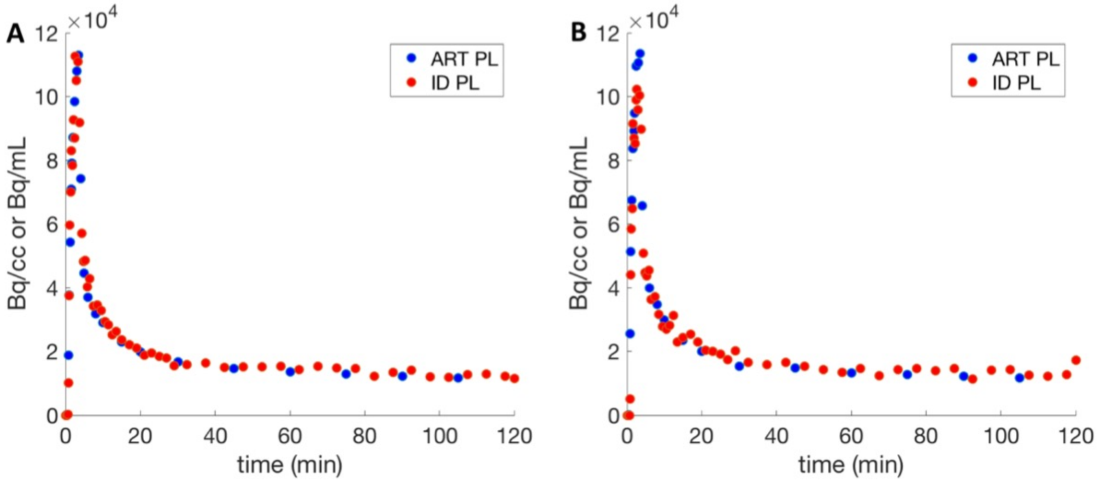


**Figure S2**: Image derived plasma curves (ID PL) obtained from the left ventricular chamber of Monkey 3 and arterial plasma curves (ART PL) obtained from blood sampling measurements plotted on the same graphs for comparison for baseline scan 1 (A) and baseline scan 2 (B).
